# Condensin-dependent chromatin condensation represses transcription globally during quiescence

**DOI:** 10.1101/320895

**Authors:** Sarah G. Swygert, Seungsoo Kim, Xiaoying Wu, Tianhong Fu, Tsung-Han Hsieh, Oliver J. Rando, Robert N. Eisenman, Jay Shendure, Jeffrey N. McKnight, Toshio Tsukiyama

## Abstract

Quiescence is a stress-resistant state in which cells reversibly exit the mitotic cell cycle and suspend most cellular processes. Quiescence is essential for stem cell maintenance and its misregulation is implicated in tumor formation. One of the conserved hallmarks of quiescent cells, from *Saccharomyces cerevisiae* to humans, is highly condensed chromatin. Here, we use Micro-C XL to map chromatin contacts at single-nucleosome resolution genome-wide to elucidate mechanisms and functions of condensed chromatin in quiescent *S. cerevisiae* cells. We describe previously uncharacterized chromatin domains on the order of 10-60 kilobases that in quiescent cells are formed by condensin-mediated chromatin loops. Conditional depletion of condensin prevents chromatin condensation during quiescence entry and leads to widespread transcriptional de-repression. We further demonstrate that condensin-dependent chromatin compaction is conserved in quiescent human fibroblasts. We propose that condensin-dependent condensation of chromatin represses transcription throughout the quiescent cell genome.

## Introduction

Quiescence is a reversible state in which cells exit the mitotic cell cycle and suspend the majority of cellular processes in response to external cues such as nutrient deprivation and contact inhibition (Cheung and Rando, 2013; Gray et al., 2004). Quiescence enables long-term cell survival and is widely conserved among eukaryotes (Allen et al., 2006; De Virgilio, 2012). For example, single-celled eukaryotes in the wild spend the majority of their time in quiescence or as spores to survive harsh conditions, and many adult mammalian tissues maintain quiescent stem cell stores (Cheung and Rando, 2013; De Virgilio, 2012; Gray et al., 2004; Valcourt et al., 2012). During quiescence entry, cells induce pathways that promote stress resistance and longevity, placing quiescence as an alternate fate to senescence and apoptosis (Cheung and Rando, 2013; Valcourt et al., 2012). Subsequent to receiving appropriate signals, quiescent cells are able to re-enter the cell cycle to resume cell division (Cheung and Rando, 2013; Gray et al., 2004; Valcourt et al., 2012). Despite its high level of conservation, the non-cycling nature of quiescent cells has made examination of quiescence difficult in any organism. However, the development of a *Saccharomyces cerevisiae* model system of quiescence has allowed for the generation of sufficient quantities of pure quiescent cells for study (Allen et al., 2006). As purified S. *cerevisiae* quiescent cells exhibit the major characteristics of mammalian quiescent cells, they are an attractive model to investigate the mechanisms underlying quiescence (Allen et al., 2006; De Virgilio, 2012; Gray et al., 2004; Li et al., 2013).

We have previously shown that budding yeast quiescent cells exhibit widespread transcriptional repression, similar to their mammalian counterparts (McKnight et al., 2015). This global decrease in gene expression is accompanied by genome-wide changes in chromatin structure, including significant increases in nucleosome occupancy and decreases in histone acetylation. Our previous work revealed that the genome-wide targeting of the histone deacetylase Rpd3 to gene promoters between stationary phase and quiescence drives this transcriptional reprogramming (McKnight et al., 2015). However, the mechanism by which these changes in chromatin structure repress transcription during quiescence is unknown. One of the highly conserved features of quiescent cells is chromatin that appears densely stained in microscopic analyses (Evertts et al., 2013; Laporte et al., 2016; Lohr and Ide, 1979; Pinon, 1978; Rawlings et al., 2011). As condensed chromatin is generally correlated with transcriptional silencing, it has been an attractive hypothesis that chromatin condensation contributes to transcriptional repression in quiescent cells (Laporte et al., 2016; Ngubo et al., 2011; Rutledge et al., 2015; Schafer et al., 2008). However, the technology to examine higher-order chromatin structure at the resolution necessary to determine its relationship to the transcription of single genes has only recently been developed (Friedman and Rando, 2015; Hsieh et al., 2015).

To investigate the role of chromatin higher-order structure in transcriptional repression during quiescence, we have employed Micro-C XL, a modified form of Hi-C capable of mapping chromatin interactions genome-wide at nucleosome-resolution (Hsieh et al., 2015; Hsieh et al., 2016). Our analyses revealed a genome-wide shift in chromatin conformation in quiescent cells. In particular, we have discovered that the condensin complex binds at many more locations throughout the quiescent cell genome than in exponentially growing cells, and induces looping between the boundaries of previously unidentified large chromatin domains. The depletion of condensin during quiescence entry prevents chromatin compaction, disrupts large chromatin domains, and de-represses genes both near boundaries and within domains genome-wide. Our results suggest that condensin represses transcription during quiescence by extruding loops that lead to chromosome condensation and boundary insulation.

## Results

### Transcriptional repression in quiescence correlates with chromatin condensation

We have previously reported, using RNA purification followed by deep sequencing (RNA-seq), that transcription is repressed genome-wide in S. *cerevisiae* quiescent cells (McKnight et al., 2015). However, recent evidence that many mRNAs are stabilized by sequestration into P-bodies during quiescence suggests that measurements of steady-state RNA levels may lead to an overestimation of transcriptional activity in quiescent cells (Aragon et al., 2006; Young et al., 2017). To measure transcriptional activity more accurately, we performed chromatin immunoprecipitation followed by deep sequencing (ChIP-seq) of RNA Polymerase II (Pol II) in exponentially growing (log) and purified quiescent cells (**Fig. S1A**). As expected, we found that Pol II enrichment is massively diminished around transcriptional start sites (TSSs) in quiescence compared to log (**Fig. S1B**). Using the peak calling algorithm incorporated into MACS software to call ChIP peaks above a 1.5-fold threshold (Zhang et al., 2008), we found that Pol II is enriched at over three times the number of genomic loci in log than in quiescence (**Fig. S1C**). Additionally, only a small subset of Pol II peaks are significantly enriched in both log and quiescent cells, consistent with massive transcriptional reprograming during quiescence entry (Roche et al., 2017).

As chromatin condensation is considered a hallmark of quiescent cells in all organisms, we sought to measure the extent of compaction in our yeast system. We used DAPI to fluorescently stain the DNA of cells synchronized in G1 and purified quiescent cells and imaged these cells by confocal microscopy **(Fig. S1D-S1E**). As expected, the chromatin of quiescent cells is significantly condensed compared to G1 cells, with the mean measured chromatin volume decreasing from 1.40 μm^3^ to 0.960 μm^3^. Together, these results indicate that widespread transcriptional repression occurs concomitantly with chromatin condensation in yeast quiescent cells. This finding led us to hypothesize that chromatin compaction may function as the mechanism by which transcription is shut-off during quiescence.

### Chromatin structure changes genome-wide between exponential growth and quiescence

In order to examine the relationship between genome-wide changes in chromatin structure and transcriptional activity between log and quiescence in detail, we performed a recently-developed variation of the Hi-C family of techniques known as Micro-C XL (Hsieh et al., 2016). During the Micro-C XL protocol, cells are crosslinked with both a short crosslinker (formaldehyde) and a long crosslinker (disuccinimidyl glutarate) and chromatin is digested down to single nucleosomes with Micrococcal nuclease. As a result, Micro-C XL is capable of mapping both short and long-range chromatin contacts genome-wide at ~150 basepair (bp) resolution (Hsieh et al., 2016).

We first generated Micro-C XL plots of log and quiescence at 5 kilobase (kb) resolution genome-wide (**Fig. 1A-B**). To ensure that differences between these experiments are reproducible, we compared biological and technical replicates using HiCRep (**Fig. S2A**) (Yang et al., 2017). The stratum-adjusted correlation coefficients between replicates averaged ~0.9, whereas those between non-replicates averaged <0.8, consistent with previous studies of reproducibility in Hi-C datasets (Yardimci et al., 2018). Therefore, we merged replicates to maximize the resolution of our analyses. By subtracting the log from the quiescence data, it is immediately apparent that long-distance interactions increase globally in both *cis* and *trans* in quiescence, as shown by a general increase in signals far from the diagonal line (**Fig. 1C**). On the other hand, very local *cis* interactions represented by dense signals along the diagonal line are favored in log. Plotting the distance decay of interactions shows interactions beyond ~ 800 bp (approximately four nucleosomes) are appreciably enriched in quiescence versus log, with log interactions falling off even more precipitously in the range of ~30 kb (**Fig. S2B**). Consistent with previous reports (Guidi et al., 2015; Laporte et al., 2016; Rutledge et al., 2015), centromere clustering decreases (**Fig. S2C**) and telomere clustering increases (**Fig. S2D**) during quiescence. Interestingly, we found a striking increase in interactions between the rDNA locus and the Chromosome XII region flanked by the centromere and the rDNA locus (termed the “pre-rDNA”) in quiescence, similar to interactions previously observed in mitotic cells (**Fig. S2E**) (Schalbetter et al., 2017). It is possible that this increase in rDNA-proximal contacts contributes to the condensation of the nucleolus seen by microscopy in yeast cells during starvation conditions (Tsang et al., 2007).

**Figure 1.**
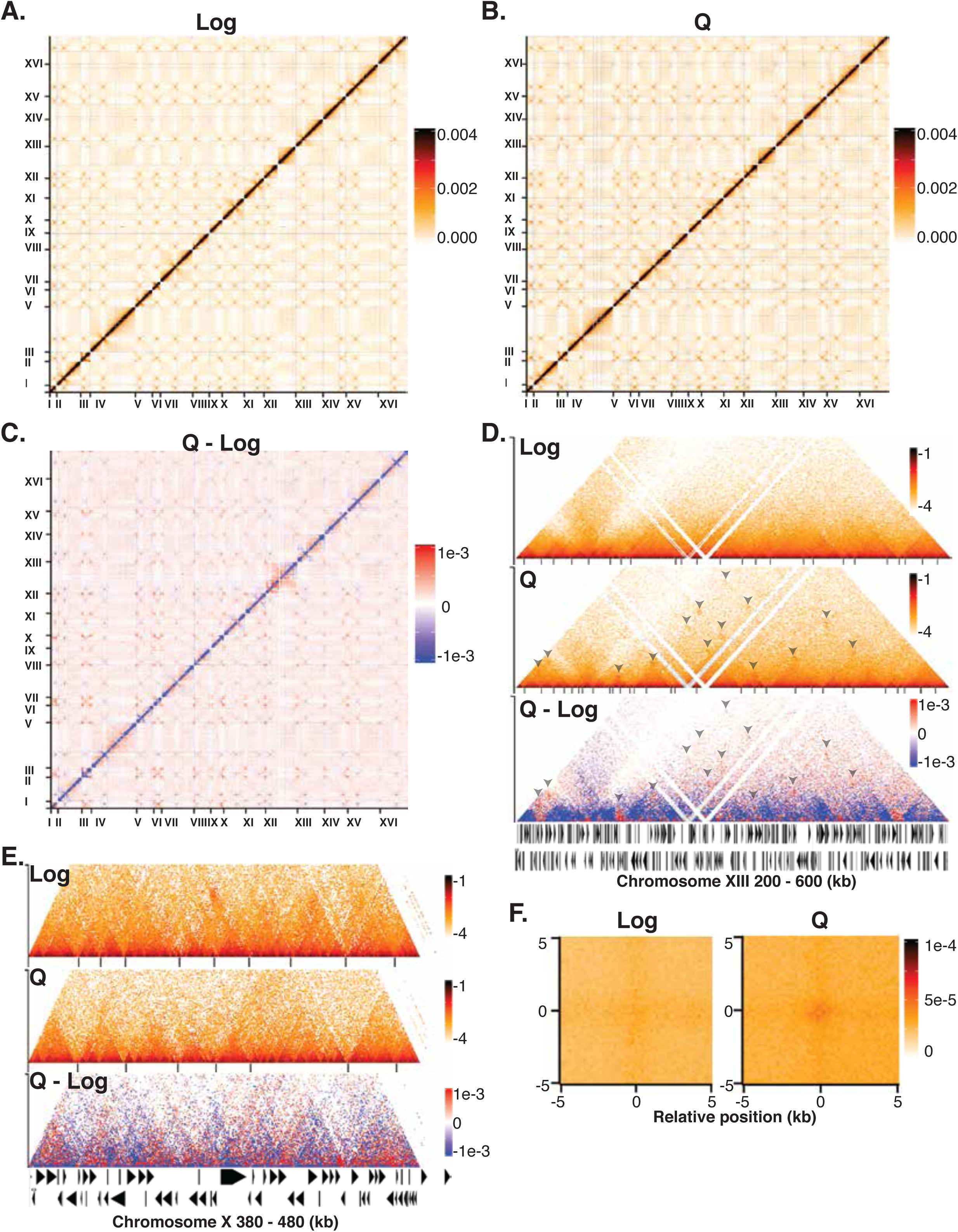
Chromatin structure changes genome-wide between exponential growth and quiescence. (**A**) Genome-wide heatmap of Micro-C XL data in exponentially growing cells (log). Each pixel represents normalized chromatin contacts between 5 kb bins. Pixels in rows and columns excluded due to insufficient read coverage are colored grey. Micro-C data here and throughout from biological and technical replicates were merged prior to normalization to read depth. (**B**) Genome-wide heatmap of Micro-C XL data in quiescent cells (Q). (**C**) Difference plot of log Micro-C data subtracted from Q. Red indicates contacts are higher in Q, blue indicates a greater number of contacts in log. (**D**) Micro-C XL data at 1 kb resolution across a region of Chromosome XIII. Dots represent normalized chromatin contacts in log scale. Black lines below contact maps indicate called boundaries of L-CIDs. Arrowheads point to dots at the corners of triangles which indicate loops formed between domain boundaries. (**E**) Micro-C XL data at 200 bp resolution across a region of Chromosome X. (**F**) Metaplots of interactions at distances of 80-320 kb between L-CID boundaries in log and Q. Pixels represent normalized interactions within 200 bp bins.

After establishing differences in long-range interactions between log and quiescence, we next generated higher-resolution interaction maps to examine the effects of quiescence entry on local interactions. The first Micro-C report identified chromosomal interaction domains (CIDs) in log budding yeast, which resemble the topologically associated domains (TADs) studied extensively in metazoans (Hsieh et al., 2015). CIDs are genomic regions on the order of one to four genes (~0.5 – 8 kb) in which chromatin interactions occur frequently. CIDs are separated by boundaries, frequently formed by highly active gene promoters, which are believed to insulate CIDs by preventing interactions across boundaries (Hsieh et al., 2015). As the number of active genes is greatly reduced and the frequency of distal chromatin interactions greatly increased in quiescence, we hypothesized that the size distribution of CIDs would increase in quiescence. However, by scanning for local minima within bin sizes consistent with CIDs, we did not observe substantial differences in CID size or number between log and quiescence (**Fig. S3A**). Instead, in genomic interaction plots at 1 kb and 200 bp resolutions, we noticed that chromatin domains on the order of 10 - 60 kb become more visible in quiescence (**Fig. 1D-1E**). Although these domains are present in log, interactions within these domains beyond ~ four nucleosomes increase in quiescence and boundaries become sharper (**Fig. 1D-1E**). Most strikingly, looping interactions between the boundaries of these domains appear frequently in quiescence (**Fig. 1D**). In fact, interactions between boundaries are apparent even at distances of 80 – 320 kb (**Fig. 1F**). Further characterization of these large domains revealed that they are of similar size and number in log and quiescence, with 709 domains with an average length of 14.3 kb in log and 806 domains with an average size of 12.7 kb in quiescence (**Fig. S3B**). Concordantly, the majority of boundaries between domains are shared between log and quiescence (**Fig. S3C**), and in both conditions boundaries occur most frequently between genes oriented in divergent and tandem directions (**Fig. S3D**) and tRNA genes (**Fig. S3E**). Although large domains on the order of ~200 kb that link regions of similar replication timing have previously been identified in budding yeast (Eser et al., 2017), these domains are of a distinct size and function. To distinguish the domains discussed in this manuscript from previously identified smaller and larger domains, we refer to them here as large CIDs (L-CIDs).

### Condensin exhibits quiescence-specific localization across the genome

In addition to their more established functions in chromatin organization during interphase and mitosis (Noma, 2017; Uhlmann, 2016; Yuen and Gerton, 2018), several previous studies have implicated Structural Maintenance of Chromatin (SMC) complexes in playing roles during quiescence. For example, temperature-sensitive condensin mutants were previously found to reduce chromatin condensation in quiescent yeast cells by microscopic analyses, and condensin is known to be essential for the compaction of the nucleolus during starvation (Laporte et al., 2016; Rutledge et al., 2015; Tsang et al., 2007; Wang et al., 2016). Intrigued by the formation of loops between large chromatin domain boundaries in quiescence, we performed a ChIP-seq screen to investigate the potential roles of SMC complexes in altering chromatin structure during quiescence. Our ChIP-seq data revealed that although cohesin and Smc5/6 ChIP peaks localized to known regions in log cells, with cohesin in particular demonstrating strong peaks of enrichment at convergent intergenic regions (Hu et al., 2011; Lengronne et al., 2004), both complexes have greatly diminished localization during quiescence (**Fig. S4A-S4B**). In contrast, the Brn1 subunit of the condensin complex showed both altered localization and a greatly increased number of peaks in quiescent cells (**Fig. 2A**). Using MACS to call enriched ChIP peaks, we determined that the number of Brn1 peaks increases from 376 in log to 1,107 in quiescence (**Fig. 2B**). Further characterization of these peaks showed that although condensin primarily binds at tRNA genes and centromeres in log, its binding shifts toward slight enrichment at tandem and strong enrichment at divergent gene promoters in quiescent cells (**Fig. 2C**). Plotting Brn1 signals alongside histone H3 and Pol II ChIP-seq signals at all TSSs ranked by Brn1 enrichment revealed that condensin is present at a small subset of highly expressed Pol II genes during log but is absent from the majority of TSSs (**Fig. 2D**). In sharp contrast, in quiescent cells, Brn1 strongly localizes to nucleosome depleted regions (NDRs) at the TSSs of the most highly expressed genes, where condensin enrichment is strongly negatively correlated with H3 enrichment (**Fig. 2E**).

**Figure 2.**
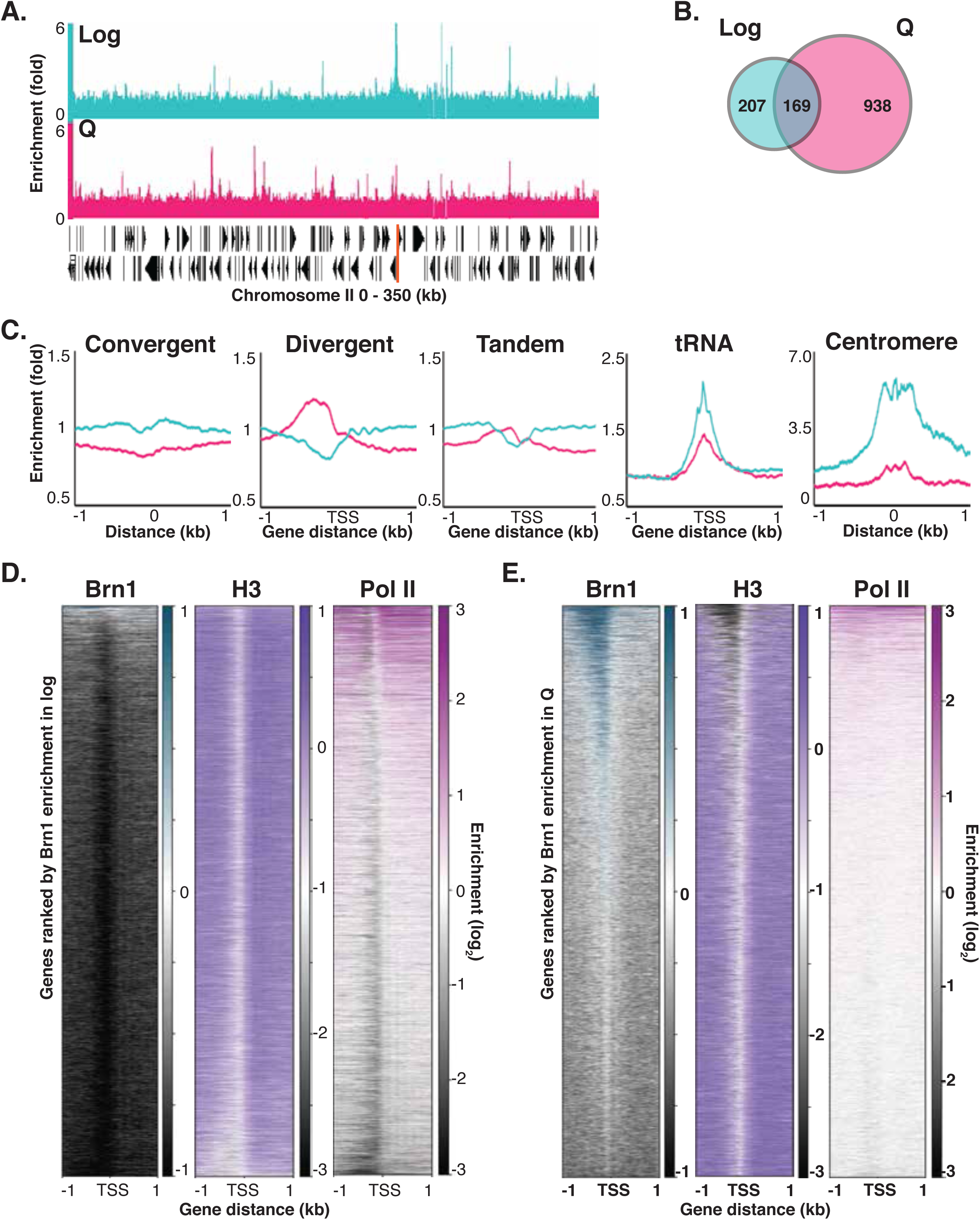
Condensin exhibits quiescence-specific localization across the genome. (**A**) Example ChIP-seq data of condensin subunit Brn1 in log and Q cells across a region of Chromosome II. The centromere is indicated by an orange line. ChIP data here and throughout were merged for biological replicates, normalized to input, and adjusted so that the genome average was set to 1-fold enrichment. (**B**) Number of Brn1 peaks above a 1.5-fold threshold called by MACS in log and Q cells. (**C**) Metaplots of average Brn1 enrichment across the indicated genomic feature in log (blue) and Q (red). (**D**) Heatmaps of ChIP-seq data for Brn1, histone H3, and RNA Polymerase II (Pol II) across all Pol II TSSs in log. All heatmaps are ranked in descending order of Brn1 enrichment in log. (**E**) Heatmaps of ChIP-seq data as in (D) in Q, ordered by strength of Brn1 enrichment in Q.

### Condensin binds the boundaries of L-CIDs during quiescence

As our ChIP data indicate that, during quiescence, condensin re-localizes to the same types of regions that form boundaries between L-CIDs, we next sought to determine whether condensin binds to L-CID boundaries in quiescent cells. Overlaying condensin ChIP data in quiescence onto Micro-C maps at 1 kb resolution demonstrates that this is true (**Fig. 3A**). Although the boundaries between L-CIDs are largely conserved between log and quiescence, only 25 percent of boundaries fall near condensin peaks in log cells, whereas 75 percent of boundaries are near condensin peaks in quiescence (**Fig. 3B, left**). Similarly, many more condensin peaks in quiescence are located near boundaries (**Fig. 3B, right**). Metaplots of Micro-C data in both log and quiescence centered on condensin peaks in quiescent cells show that the location of condensin in quiescence predicts domain boundaries in both log and quiescence (**Fig. 3C**). In fact, approximately 75 percent of domain boundaries in log and quiescence fall within 1 kb of a condensin peak in quiescence (**Fig. 3D**). However, in quiescent cells, condensin binding coincides with a slight increase in longer-range *cis* interactions within domains and a slight decrease in interactions across boundaries (**Fig. 3C**). Consistently, condensin binding in quiescence correlates with the strength of boundary insulation, suggesting that condensin itself may be strengthening boundaries in quiescent cells (**Fig. 3E**). In addition, condensin binding sites near boundaries in quiescence insulate chromatin interactions more robustly in quiescence than the same locations in log (**Fig. S3F**). Curiously, despite the correlation between strong boundaries and condensin binding in quiescent cells, the majority of boundaries in log are depleted not only in condensin but also in cohesin and Smc5/6 (**Fig. S4C-D**). Finally, to examine the possibility that condensin is responsible for the long-range interactions observed between boundaries of L-CIDs in quiescent cells, we plotted interactions between condensin peaks located at boundaries on the same chromosome (**Fig. 3F**). Quiescent but not log cells demonstrated highly localized interactions between condensin peaks. These results collectively support a model in which condensin is involved in forming loops between L-CID boundaries during quiescence.

**Figure 3.**
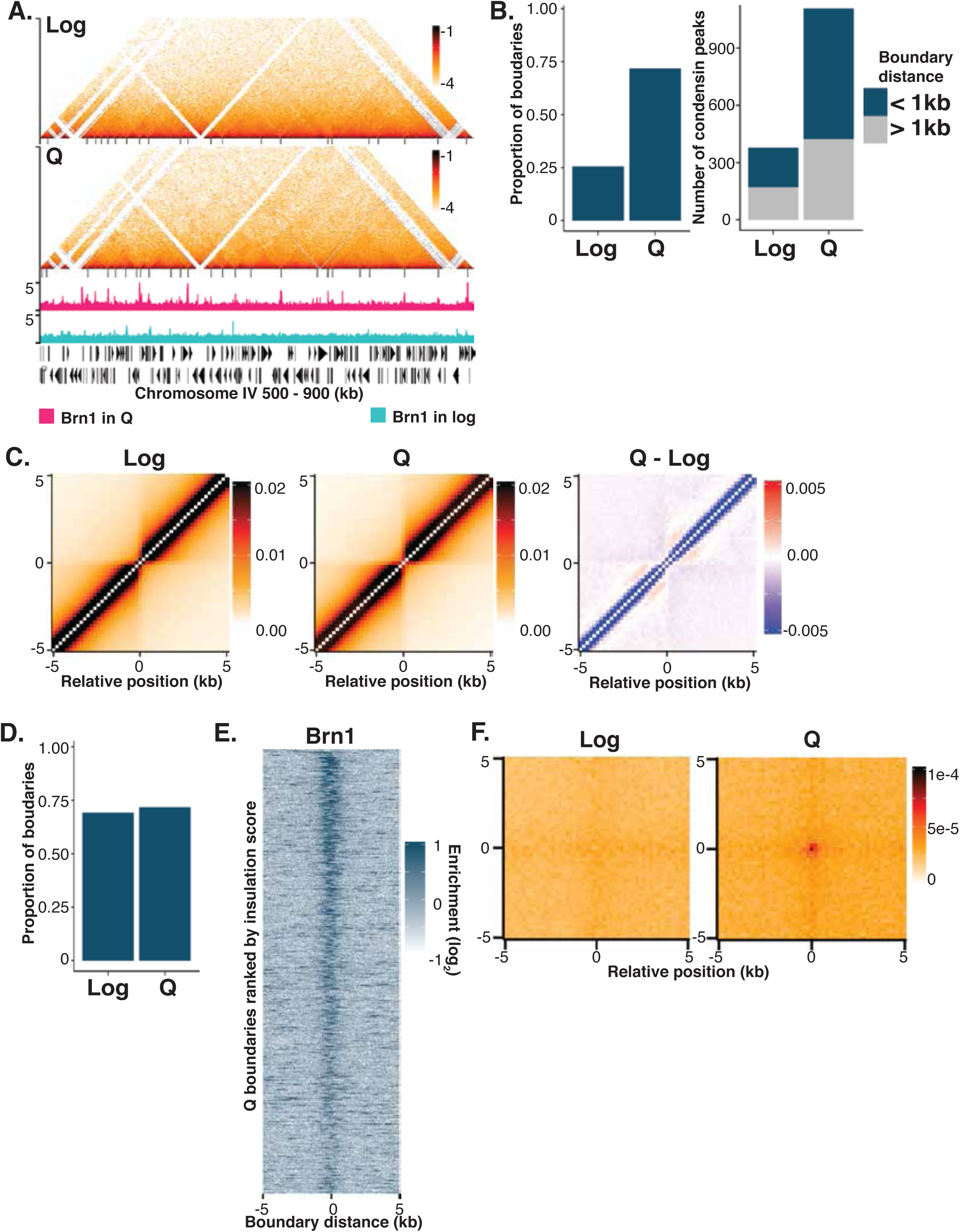
Condensin binds the boundaries of L-CIDs. (**A**) Micro-C XL contact maps at 1 kb resolution across a region of Chromosome IV. Called L-CID boundaries are indicated by black lines below contact maps. ChIP-seq data of Brn1 in Q (red) and log (blue) are aligned beneath contact maps. (**B**) Left: Proportion of L-CID boundaries called in log and Q within 1 kb of a Brn1 peak in log and Q, respectively. Right: Number of Brn1 peaks in log and Q within 1 kb (in blue) and more than 1 kb away (grey) from a called L-CID boundary in log and Q, respectively. (**C**) Metaplots of average Micro-C XL contacts at 200 bp resolution in log, Q, and log subtracted from Q centered on Brn1 peaks in Q. Pixels represent normalized contacts. In the subtraction plot, red represents contacts that are higher in Q and blue represents contacts that are higher in log. (**D**) Proportion of L-CID boundaries in log and Q within 1 kb of a Brn1 peak in Q. (**E**) Heatmap of Brn1 enrichment at called L-CID boundaries in Q. Boundaries are ranked in descending order by strength of insulation. (**F**) Metaplots of average Micro-C contacts at 200bp resolution in log and Q between condensin peaks located at L-CID boundaries in Q. The plot shows contacts at distances of 80 – 320 kb.

### Condensin is required for chromatin condensation and L-CID formation in quiescence

The binding of condensin to L-CID boundaries in quiescence suggested that condensin may play a role in chromatin condensation in quiescence. To test this, we constructed doxycycline-inducible tet-off depletion strains of condensin subunit Smc4 (tet-*SMC4*). By adding doxycycline to cells just prior to the diauxic shift and replenishing it regularly throughout quiescence entry, we are able to reproducibly achieve >90% depletion of the Smc4 protein (**Fig. S5A**). Consistent with our hypothesis that condensin compacts chromatin in quiescence, DAPI staining and confocal microscopy of tet-*SMC4* quiescent cells formed in the presence (*SMC4*-off) or absence (*SMC4*-on) of doxycycline demonstrates a significant increase in chromatin volume from 1.04 μm^3^ to 1.29 μm^3^ upon condensin depletion (**Fig. 4A-4B**). Addition of doxycycline during quiescence entry to control cells does not alter chromatin volume (**Fig. S5B-S5C**). *SMC4*-on quiescent cell chromatin is slightly decondensed as compared to wild-type (WT) (**Fig. S1E**), suggesting that the epitope tagging of Smc4 and/or the expression of the *SMC4* gene via a synthetic promoter leads to a slight loss of function of condensin. We therefore focused on comparing data from *SMC4*-off quiescent cells to WT quiescent cells to elucidate condensin functions in this manuscript, with *SMC4*-on data presented in supplementary figures.

**Figure 4.**
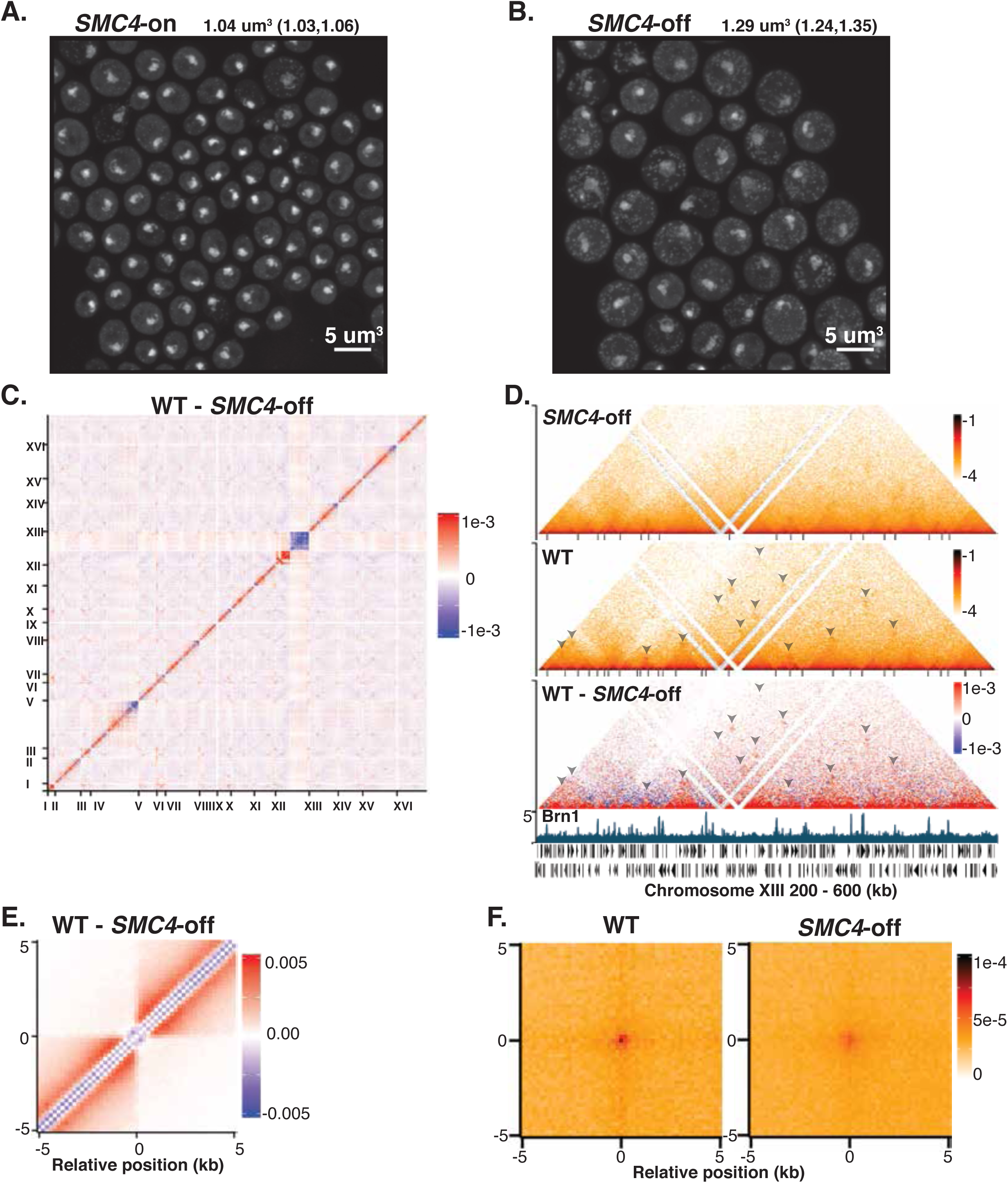
Condensin is required for chromatin condensation and L-CID formation in quiescence. (**A-B**) Microscopy of Q cells purified from Smc4-tet-off strains in the absence (*SMC4*-on) or presence (*SMC4*-off) of doxycycline. DNA is stained with DAPI. Numbers shown are the mean volume of stained chromatin and the lower and upper bounds of the 95% confidence interval. (**C**) Genome-wide Micro-C XL contact map at 5 kb resolution of *SMC4*-off data subtracted from wild-type (WT) Q data. Red indicates interactions are higher in WT Q, blue indicates interactions are higher in *SMC4*-off. (**D**) Micro-C XL data at 1 kb resolution across a region of Chromosome XIII. Called L-CID boundaries are indicated by black lines beneath contact maps. Arrowheads point to looping interactions between domain boundaries. Brn1 ChIP-seq data from WT Q cells are aligned below. (**E**) Metaplot of the differences in Micro-C signals at 200 bp resolution between *SMC4*-off and WT Q centered on Brn1 peaks in Q. (**F**) Metaplots of Micro-C signals at 200 bp resolution in WT Q and *SMC4*-off cells between condensin peaks located at boundaries in Q at distances of 80 – 320 kb.

To examine the effect of condensin depletion on chromatin contacts in detail, we performed Micro-C XL in *SMC4*-off (**Fig. S6A**) and *SMC4*-on (**Fig. S6B-S6C**) quiescent cells. Subtracting *SMC4*-off from WT data genome-wide at 5 kb resolution indicates that long-range *trans* interactions are increased, as shown by increase in blue signals away from the diagonal line. In contrast, short-range local interactions are sharply decreased in *SMC4*-off cells, as represented by red signals along the diagonal line (**Fig. 4C**). In particular, condensin depletion leads to a loss of interactions in *cis* on the scale of ~1-50 kb (**Fig. S2B**). *SMC4*-off cells also display an increase in centromere clustering and a slight decrease in telomere clustering compared to WT, although *trans* interactions between regions near centromeres decrease (**Fig. S2C-S2D**). Condensin depletion also diminished interactions between the centromere-distal region of Chromosome XII and the rDNA, consistent with a role for condensin in compaction of the pre-rRNA (**Fig. 4C** and **S2E**) (Schalbetter et al., 2017). Although condensin-mediated interactions between tRNA genes and the rDNA have been seen to be lost during nutrient depletion (Belagal et al., 2016; Haeusler et al., 2008), these interactions did not appear to change appreciably between log and quiescence or between WT and *SMC4*-off quiescent cells (**Fig. S2G**). Curiously, *SMC4*-off cells do display a striking increase in the isolation of many telomeres and entire rDNA-distal portion of the right arm of Chromosome XII, reflected by a loss of *trans* contacts between these regions and the rest of the genome (**Fig. 4C, S2F**). These results may be caused by a major alteration in nuclear organization due to changes in centromere, telomere, and nucleolar location and condensation.

We next examined the *SMC4*-off Micro-C data at high resolution. As expected, condensin depletion did not affect the size or number of CIDs (**Fig. S2A**). However, plotting data at 1 kb resolution showed marked alteration to the structure of L-CIDs leading to the loss of interactions within L-CIDs as well as a decrease in looping interactions between L-CID boundaries (**Fig. 4D**). To demonstrate these results more clearly, we subtracted *SMC4*-off from WT Micro-C data at 200 bp resolution and plotted the average data genome-wide centered at condensin ChIP-seq peaks in quiescence (**Fig. 4E**). These data show a global decrease in interactions within L-CIDs upon condensin depletion, and an increase in interactions across sites of condensin peaks where boundaries occur. In addition to a decrease in interactions across condensin peaks, condensin depletion also dramatically reduced the number of L-CIDs that could be called by our boundary-locating algorithm from 806 to 492 (**Fig. S3B**), supporting a role for condensin as an insulator at L-CID boundaries during quiescence. Insulation scores around quiescent Brn1 peaks also sharply increased in *SMC4*-off cells, indicating a decrease in insulation at condensin binding sites (**Fig. S3F**). Finally, we also plotted long-range *trans* interactions centered at condensin boundary peaks in quiescence (**Fig. 4F**). Consistent with the analysis at individual loci (**Fig. 4D**), global interactions between condensin binding sites at boundaries decrease when condensin is depleted. Interactions between condensin binding sites do not entirely disappear in *SMC4*-off, however, which is likely due to our inability to deplete condensin completely in these cells (**Fig. S5A**). However, we cannot rule out the presence of an ancillary factor that assists in loop formation at condensin binding sites. Cumulatively, the role of condensin in mediating long-range interactions, insulating domains, and increasing intra-domain interactions supports our model that condensin mediates chromatin compaction in quiescent cells by forming loops between L-CID boundaries.

### Condensin is required for transcriptional repression during quiescence

The establishment of a system in which chromatin condensation in quiescent cells can be conditionally disrupted by the inducible depletion of condensin allowed us to test our hypothesis that chromatin compaction represses transcription in quiescence. To this end, we performed Pol II ChIP-seq in purified quiescent *SMC4*-off cells and compared this to our Micro-C XL data (**Fig. 5A-5B**). Our results indicate that condensin depletion leads to widespread gene de-repression in quiescence, with the number of Pol II ChIP-seq peaks increasing from 864 in WT cells to 2,021 in *SMC4*-off (**Fig. 5C**). In some cases, condensin binds at the intergenic regions between divergent genes where only one of the two genes is highly transcribed, forming a boundary between two L-CIDs (**Fig. 5A-5B**). On average, condensin depletion leads to increased transcription of genes on both sides of condensin binding sites (**Fig. 5D**). Heatmaps of WT Q and *SMC4*-off Pol II data ordered by strength of condensin signals show that the largest increases in Pol II occupancy in *SMC4*-off cells occur at genes with Brn1 bound to strong NDRs in their promoters (**Fig. 2E, 5E**). However, many genes not bound by Brn1 were also de-repressed in *SMC4*-off cells (**Fig. 5A-5B, 5E**).

**Figure 5.**
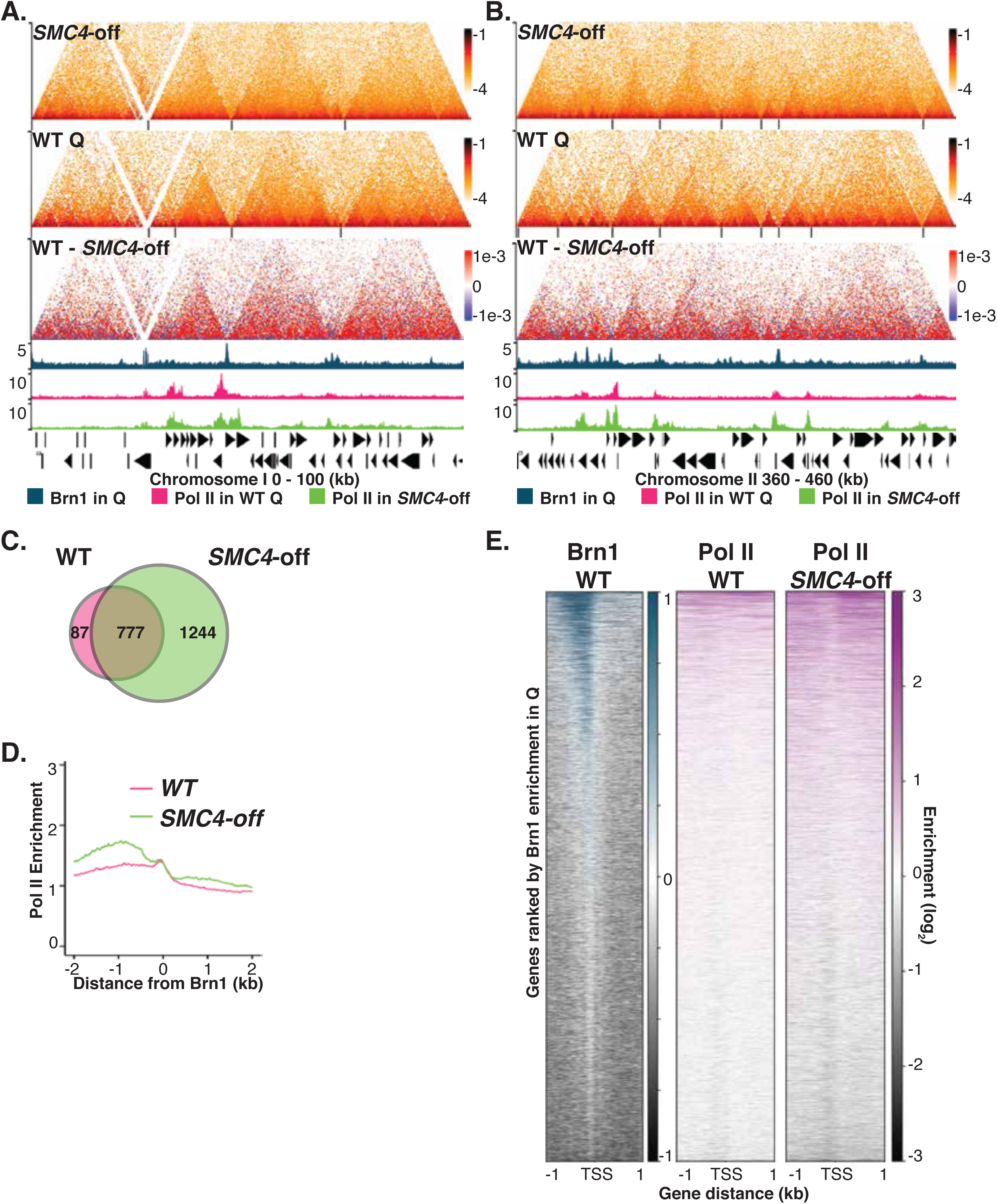
Condensin is required for transcriptional repression during quiescence. (**A-B**) Micro-C XL data in *SMC4*-off and WT Q at 200 bp resolution across a region of Chromosome I (A) and a region of Chromosome II (B). L-CID boundaries are marked by black lines beneath contact maps. ChlP-seq data of Brn1 binding in WT Q (blue), Pol II binding in WT Q (red), and Pol II binding in *SMC4*-off (green) are aligned below. (**C**) The number of Pol II ChIP-seq peaks above a 1.5-fold threshold called by MACS software in WT Q and *SMC4*-off. (**D**) Metaplot of Pol II enrichment over Brn1 peaks in WT Q. The plot is oriented such that the higher levels of Pol II signals in WT Q are on the left side of the average Brn1 peak. (**E**) Heatmaps of Brn1 binding in WT Q and Pol II binding in WT Q and *SMC4*-off at all Pol II TSSs. All heatmaps are ranked in descending order of Brn1 enrichment in WT Q.

### Condensin represses transcription globally during quiescence by condensing chromatin

The fact that condensin depletion leads to the de-repression of genes located away from Brn1 peaks suggests that condensin can repress genes independently from its direct actions at promoters. To investigate this possibility further, we examined our Micro-C and ChIP-seq data and found that as expected, many genes located well within domains away from condensin sites at L-CID boundaries gain Pol II in *SMC4*-off cells (**Fig. 6A**). Although a majority of the genes that become most de-repressed in *SMC4*-off are located within a few hundred bp of a condensin peak, a significant number are well over 1 kb away from the nearest Brn1 peak (**Fig. 6B**). Additionally, a subset of genes immediately adjacent to condensin binding sites do not become de-repressed when condensin is depleted, even though boundaries between nearby L-CIDs are lost and the surrounding chromatin becomes decondensed (**Fig. 6A-6B**). These results suggest: 1) that condensin mediates the transcriptional repression of genes nearby and at a distance by compacting chromatin, and 2) that chromatin condensation in quiescence is a cause and not a consequence of transcriptional repression.

**Figure 6.**
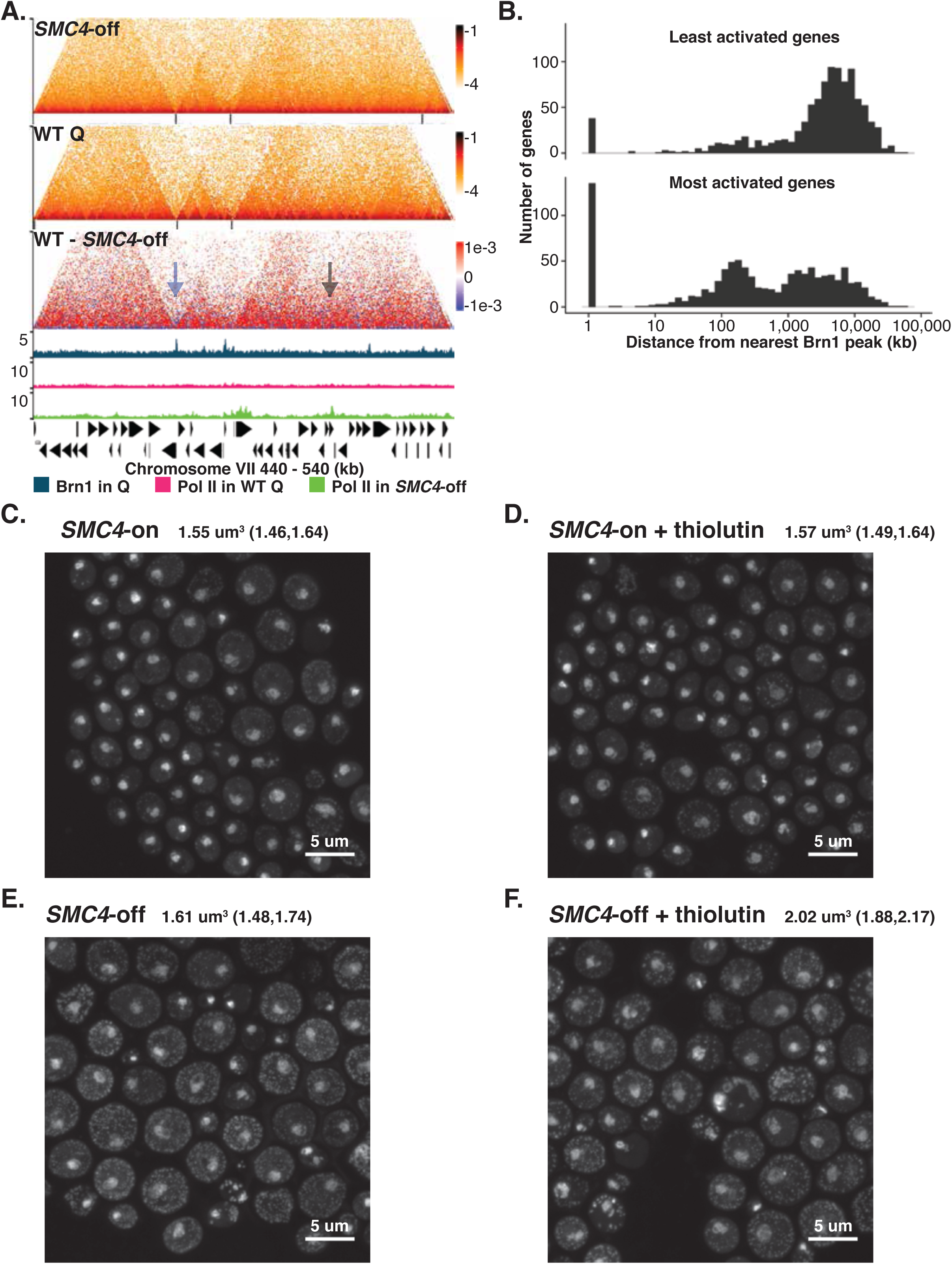
Condensin represses transcription globally by condensing chromatin. (**A**) Micro-C XL data in *SMC4*-off and WT Q at 200 bp resolution across a region of Chromosome VII. L-CID boundaries are marked by black lines beneath contact maps. ChIP-seq data of Brn1 binding in WT Q (blue) Pol II binding in WT Q (red), and Pol II binding in *SMC4*-off (green) are aligned below. Blue arrow points to a locus where a prominent Brn1 ChIP-seq peak does not align with an increase in Pol II signals in *SMC4*-off. Grey arrow points to a locus where an increase in Pol II signals in *SMC4*-off does not overlap with a Brn1 peak or a large domain boundary in WT Q. (**B**) Histograms of the number of genes with TSSs within a given distance of Brn1 ChIP-seq peaks in WT Q. Top shows the 1,000 genes with the lowest ratio of Pol II signals in *SMC4*-off over WT Q. Bottom shows the 1,000 genes with the highest ratio of Pol II signals in *SMC4*-off over WT Q. (**C-F**) Microscopy of a mixed population of Smc4-tet-off cells from a 4-day culture in the absence (*SMC4*-on) or presence (*SMC4*-off) of doxycycline with or without the addition of thiolutin. DNA is stained with DAPI. Numbers shown are the mean volume of stained chromatin and the lower and upper bounds of the 95% confidence interval.

To further test the causality relationship between chromatin decompaction and transcriptional de-repression, we performed DAPI staining and microscopy in WT and tet-*SMC4* cells in the presence of thiolutin, an RNA polymerase inhibitor (**Fig. 6C-6F**). Thiolutin was added at the diauxic shift before the differences in chromatin size in *SMC4*-on and *SMC4*-off cells becomes apparent, and was replenished every twelve hours during quiescence entry. As the addition of thiolutin during quiescence entry leads to the death of the majority of cells prior to day seven, the time at which we normally purify quiescent from non-quiescent cells, we examined mixed populations of cells from four-day cultures. On day four, the presence of thiolutin in *SMC4*-on cultures did not lead to a change in chromatin volume (**Fig. 6C-6D**). However, if persistent transcription were the cause of the lack of chromatin condensation upon condensin depletion, then the addition of thiolutin would be expected to restore chromatin condensation in *SMC4*-off cells. Instead, thiolutin addition to *SMC4*-off cells even further decondenses chromatin (**Fig. 6E-6F**). These data show that transcription is not the cause of chromatin decompaction upon condensin depletion, and supports the notion that chromatin compaction is the cause, not the result of transcriptional repression.

### Condesin-dependent chromatin condensation is a conserved feature of quiescent cells

Having established condensin-mediated chromatin condensation as a feature of quiescent budding yeast cells, we sought to determine if a similar mechanism might be conserved during human cell quiescence. To this end, we cultured human foreskin fibroblasts (HFFs) in low serum media for four days to induce quiescence. In order to examine chromatin condensation between growth and quiescence, we performed DAPI staining and immunofluorescence on G1 and quiescent cells (**Fig. 7A-7B**). Only cyclin D1 positive G1 cells and ki-67 negative quiescent cells were analyzed. We observed a statistically significant decrease in the overall nuclear area of quiescent cells, from 269 to 162 μm^2^, consistent with an increase in chromatin condensation during quiescence (**Fig. 7D**). Interestingly, we also observed the formation of discrete, densely staining puncta in quiescent cells that are present at only low levels in G1 (**Fig. 7A-7B, 7E**). We next cultured cells to quiescence transfected with either scrambled control small inhibitory RNA (siRNA) or siRNA targeting Smc2, a subunit of both the human condensin I and condensin II complexes. Transfection with targeted siRNAs leads to the knockdown of Smc2 to approximately 10 percent of the level of protein present in cells transfected with control siRNA (**Fig. S7A**). Consistent with our data from *SMC4*-off budding yeast cells, we found that knockdown of Smc2 prevents chromatin compaction during HFF quiescence entry, with DAPI staining leading to a mean nuclear area of 350 μm^2^ (**Fig. 7C-7D**). Smc2 knockdown also leads to a significant loss of the densely staining puncta that form in quiescent cells transfected with control siRNA (**Fig. 7C, 7E**). Collectively, these data show that the role of condensin in chromatin condensation during quiescence is conserved in human fibroblast cells.

**Figure 7.**
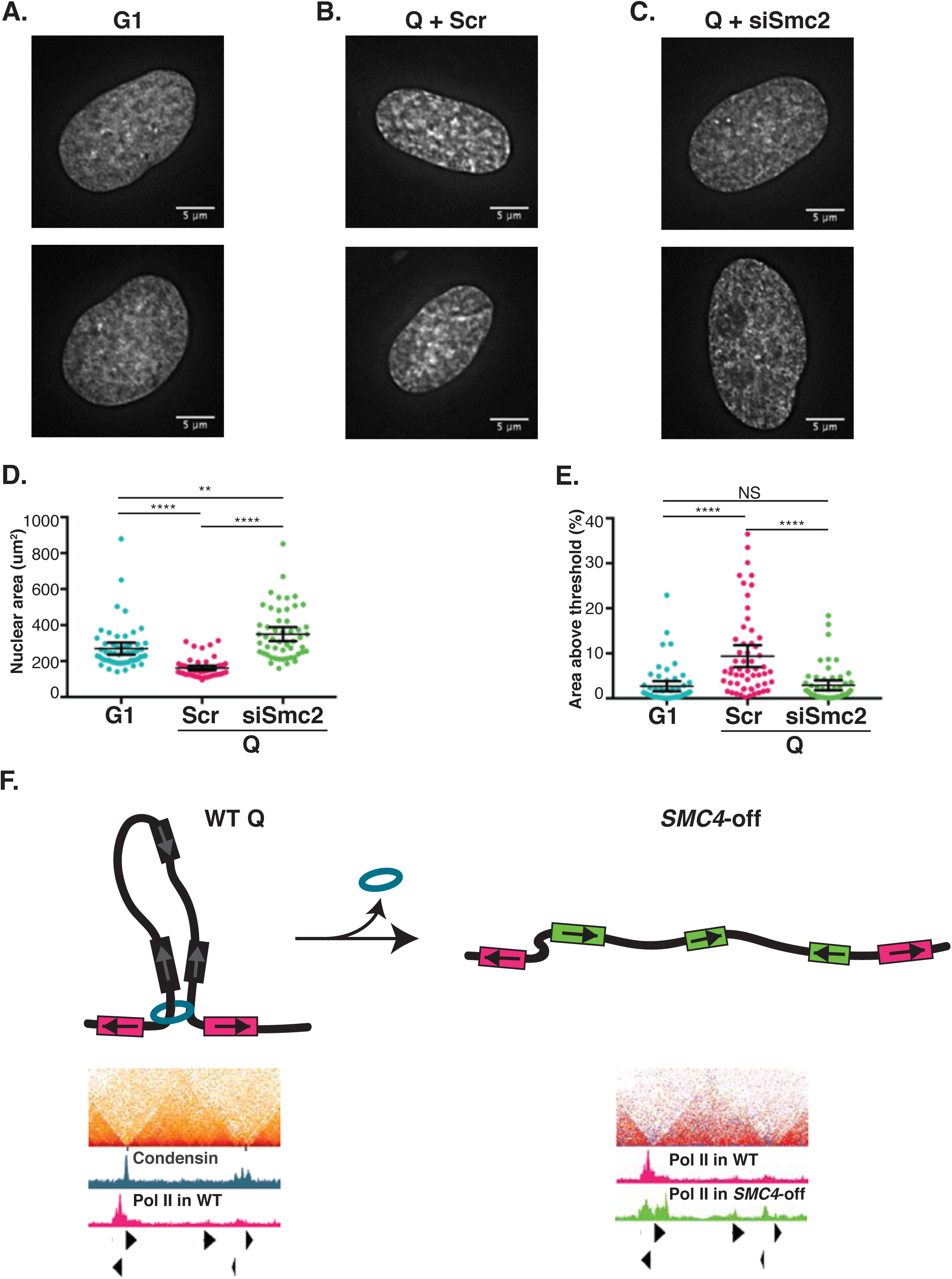
Condensin-dependent chromatin condensation is a conserved feature of quiescent cells. (**A**) DAPI staining of HFF cells in G1. (**B**) DAPI staining of quiescent HFF cells transfected with scrambled control siRNA. (**C**) DAPI staining of quiescent HFF cells transfected with siRNA targeting Smc2. (**D**) Measured nuclear area in G1 cells and quiescent cells transfected with scrambled control siRNA (Scr) or siRNA targeting SMC2. Large bars indicate the mean, small bars the upper and lower bounds of the 95% confidence interval. Bars above with asterisks indicate statistical significance. (**E**) To quantify differences in DAPI puncta, we measured the area of staining above the background threshold in each condition. Large bars indicate the mean, small bars the upper and lower bounds of the 95% confidence interval. Bars above with asterisks indicate statistical significance. NS indicates not significant. (**F**) A model for condensin function in quiescent cells. Condensin (blue) binds to nucleosome depleted regions at the promoters of Pol II-occupied divergent genes (red boxes) in Q cells. The binding of condensin forms or reinforces a loop linking L-CID boundaries. Increased proximity of chromatin within the extruded loop results in an increase in intra-domain interactions. Condensin represses the transcription of genes (black boxes) through increased chromatin compaction and insulation. When condensin is depleted during quiescence entry, L-CID formation is disrupted, looping between boundaries decreases, chromatin within L-CIDs fails to condense, and interactions across boundary regions increase. This leads to failed transcriptional repression of genes (green boxes) due to the lack of compaction and insulation.

## Discussion

In this report, we identified L-CIDs, chromatin domains in S. *cerevisiae* that are much larger than the previously characterized CIDs, and found that condensin binds to the boundaries of these domains in quiescent cells. Condensin plays key roles in the structure of L-CIDs in quiescence, both by creating L-CID boundaries, and by increasing interactions within L-CIDs that lead to the condensation of chromatin within loops. Importantly, actions by condensin lead to transcriptional repression, revealing a previously unknown mechanism for global transcriptional repression in quiescent cells.

One of our unexpected findings is that although looping interactions between L-CID boundaries are not present in log, the locations of L-CID boundaries are largely shared between log and quiescent cells. Even more puzzling is the fact that neither condensin, cohesin, nor Smc5/6 are located at the majority of L-CID boundaries in log, whereas condensin depletion leads to the loss of nearly half of all boundaries in quiescence. This preservation of L-CID boundaries between log and quiescent cells resembles that of metazoan TADs, where the strongest boundaries persist throughout cell types and are even preserved across species (Barutcu et al., 2018; Dixon et al., 2015; Dixon et al., 2012). We suspect that additional factors are involved in the formation of boundaries, especially in log cells. Previous studies in a broad range of eukaryotes from fission yeast to human cells have discovered that sequence-specific transcription factors are responsible for recruiting condensin to specific loci during mitosis and differentiation (D’Ambrosio et al., 2008; Haeusler et al., 2008; Kim et al., 2016; Nakazawa et al., 2008; Robellet et al., 2017; Yuen et al., 2017). It is possible that quiescence-specific transcription factors similarly directly recruit condensin to active promoters during quiescence. Alternately, transcription factors bound at actively transcribed genes or even divergent gene promoters themselves may facilitate the creation of chromatin boundaries by acting as barriers to extruding chromatin loops.

Although condensin has previously been linked to the formation of large chromatin domains in *Schizosaccharomyces pombe* and metazoan cells, the biological roles for chromatin domain formation have not been well understood (Kakui et al., 2017; Kim et al., 2016; Tanizawa et al., 2017; Yuen and Gerton, 2018). It is especially noteworthy that, although L-CIDs in quiescent budding yeast resemble metazoan TADs, the functions of these large domains vary widely. While condensin binding has been associated with mediating interactions between strongly insulating TAD boundaries at some locations in mouse embryonic stem cells, removal of condensin leads to the deactivation of nearby genes (Yuen et al., 2017). Recent Hi-C analyses in immortalized human colorectal cancer cells in which cohesin was conditionally depleted revealed that the loss of cohesin-dependent domains led to transcriptional changes, both activating and repressing, of about 300 genes, or slightly over one percent of all genes in the human genome (Rao et al., 2014). This is consistent with the proposed role of cohesin-loops mediating contacts between gene promoters and enhancers while restricting aberrant promoter/enhancer contacts. In contrast, long-range acting enhancers are not conserved in budding yeast (Bulger and Groudine, 2011), and yet depletion of condensin in quiescent budding yeast cells led to the transcriptional activation of approximately twenty percent of all genes. Although condensin is known to mediate the repression of tRNA-proximal Pol II genes by clustering tRNA genes near the nucleolus in log yeast cells (Haeusler et al., 2008), condensin binding is reduced at tRNA genes in quiescence and tRNA/rDNA contacts are not strongly enriched in quiescent yeast cells. Interestingly, the roles of condensin in quiescent yeast cells closely resemble those of the condensin dosage compensation complex (DCC) responsible for X-chromosome inactivation in *Caenorhabditis elegans*. The DCC is known to bind strongly insulating TAD boundaries on the inactive X-chromosome, and preventing DCC binding leads to transcriptional depression both near and away from the DCC site through an unknown mechanism (Crane et al., 2015). Similarly, our results suggest that the formation of L-CIDs during quiescence represses transcription through a previously unknown, globally acting mechanism.

How might condensin repress transcription so broadly in quiescent cells? Our data suggest that, during quiescence entry, the condensin complex targets NDRs at promoters of actively transcribed genes and facilitates boundary formation between L-CIDs (**Fig. 7F**). We envision that condensin alters chromatin interactions to repress transcription via at least two mechanisms. First, the extrusion of loops between L-CID boundaries increases chromatin contacts within loops. This chromatin compaction likely represses transcription by sterically blocking the accessibility of DNA to transcription factors and Pol II. Second, at many divergent intergenic regions where one gene is more highly transcribed, condensin binding between the two genes may additionally prevent the aberrant activation of normally silent or weakly expressed genes across condensin binding sites, potentially by blocking the effects of transcriptional activators at the promoters of active genes through boundary insulation. Although it has previously been difficult to determine whether chromatin condensation is a cause or effect of transcriptional repression, we have shown that even in the presence of thiolutin, an RNA polymerase inhibitor, condensin depletion leads to the decondensation of chromatin in quiescent cells. These results strongly support the hypothesis that chromatin compaction is the cause of transcriptional repression in quiescent cells.

A previous study used transmission electron microscopy to discover that quiescent naïve mouse T-cells have densely compacted chromatin, and that this chromatin condensation is lost upon T-cell activation (Rawlings et al., 2011). Naïve T-cells from mice harboring a mutation in the kleisin-ß subunit of condensin II do not condense their chromatin and fail to silence genes that are only expressed upon T-cell activation in WT cells (Rawlings et al., 2011). Our data show that quiescent human fibroblasts display puncta of highly condensed chromatin and decreased overall nuclear areas, and that this chromatin condensation similarly requires condensin. In sum, these results support the possibility that condensin-mediated chromatin condensation is a conserved mechanism during quiescence.

## Acknowledgements

The authors would like to thank members of the Tsukiyama and D. Koshland labs for insightful input and discussion, and in particular Samuel Cutler for advice on data analysis. Research was supported by grants F32GM120962 and T32CA009657 for S.G.S., R01GM079205 for O.J.R., RO1CA057138 for R.N.E., and R01GM111428 for T.T. J.S. is an investigator of the Howard Hughes Medical Institute. Fred Hutchinson Cancer Research Center Shared Resources were supported by the Fred Hutch/University of Washington Cancer Consortium (P30 CA015704).

## Author Contributions

S.G.S., J.N.M., and T.T. designed the experiments. S.G.S. performed all genomic experiments. S.G.S. and S.K. analyzed the ChIP-seq data. S.K. analyzed the Micro-C XL data with supervision from J.S. X.W. performed experiments in human cells with the supervision of B.N.E. T.F. performed microscopy experiments in yeast and assisted in cell preparation. T.-H.H. and O.J.R. provided the Micro-C protocol in advance of its publication, and T.-H.H. oversaw training for S.G.S. in the Micro-C method. S.G.S. and T.T. prepared the manuscript, and all authors assisted in editing of the manuscript. T.T. supervised S.G.S., T.F., and J.N.M. Fred Hutchinson Cancer Research Center Shared Resources were supported by the Fred Hutch/University of Washington Cancer Consortium (P30 CA015704).

---

**Figure S1.**
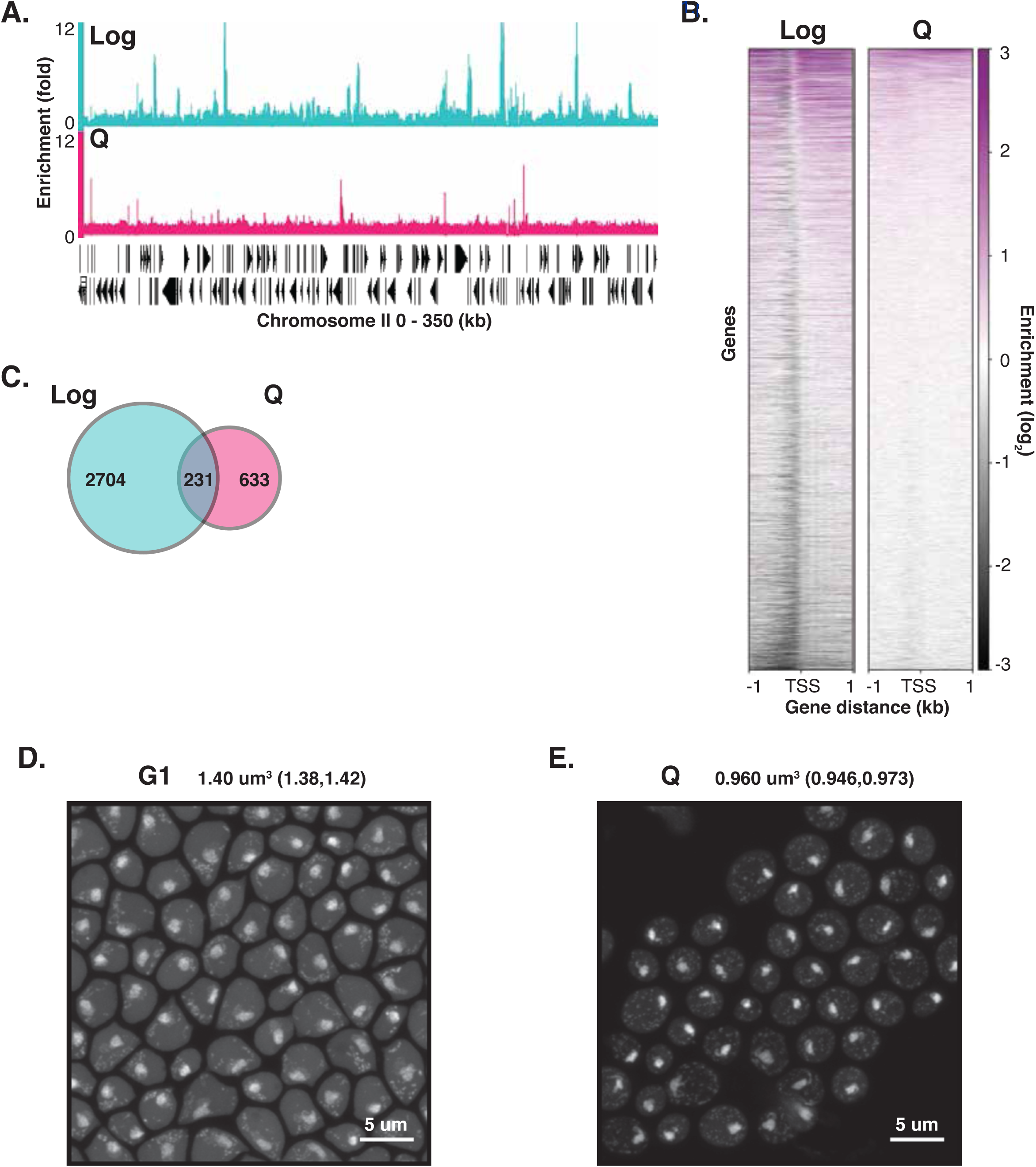
Transcriptional repression in quiescence correlates with chromatin condensation. (**A**) Example ChIP-seq of Pol II in log and Q cells across a region of Chromosome II. (**B**) Heatmaps of Pol II signals across all Pol II TSSs in log and Q. Genes are ranked in descending order of Pol II occupancy in each condition. (**C**) The number of Pol II ChIP-seq peaks above a 1.5-fold threshold called by MACS software. (**D-E**) Microscopy of cells arrested in G1 and purified Q cells. DNA is stained with DAPI. Numbers shown are the mean volume of stained chromatin and the lower and upper bounds of the 95% confidence interval.

**Figure S2.**
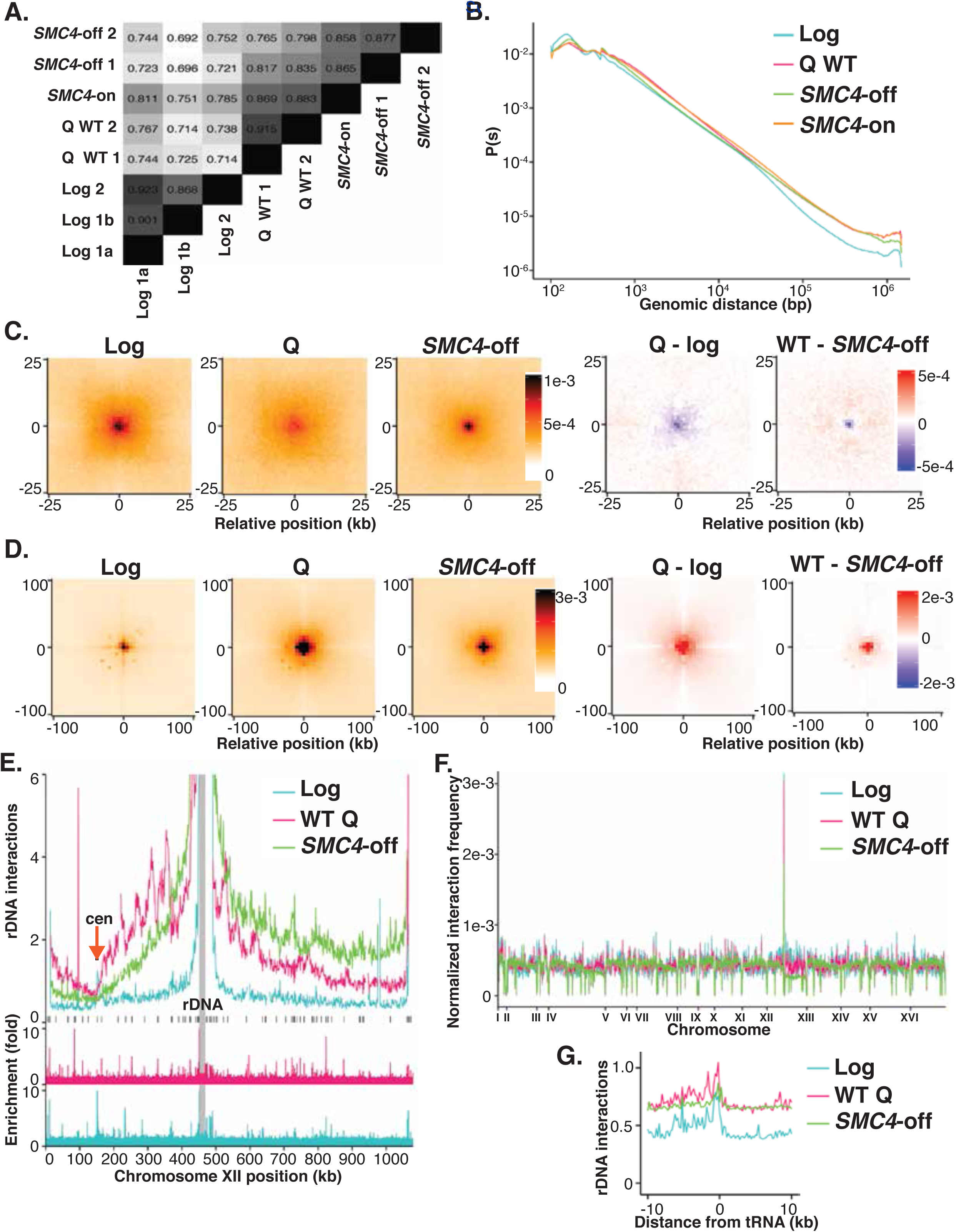
Chromatin conformation changes globally between log, WT quiescent, and *SMC4*-off quiescent cells. (**A**) Stratum adjusted correlation coefficients as determined by the HiCRep package for all Micro-C XL experiments. Biological replicates are indicated by number, technical replicate indicated by letter. Shading is scaled to degree of similarity between samples. (**B**) Distance decay plot of Micro-C XL signals showing normalized contacts across distances of 10^2^ − 1.5×10^6^ bp. (**C-D**) Metaplots of average Micro-C XL signals at 200 bp resolution of contacts between pairs of centromeres (C) and telomeres (D). (**E**) Plot of intra-chromosomal interactions between the rDNA locus and Chromosome XII detected by Micro-C. L-CID boundaries in WT Q are marked beneath by black lines, the centromere and rDNA are labeled. ChIP-seq data of Brn1 in WT Q (red) and log (blue) are aligned below. (**F**) Normalized total interaction frequency across all genomic positions in 5 kb bins genome-wide. (**G**) Metaplot of average Micro-C interactions between tRNA genes and the rDNA.

**Figure S3.**
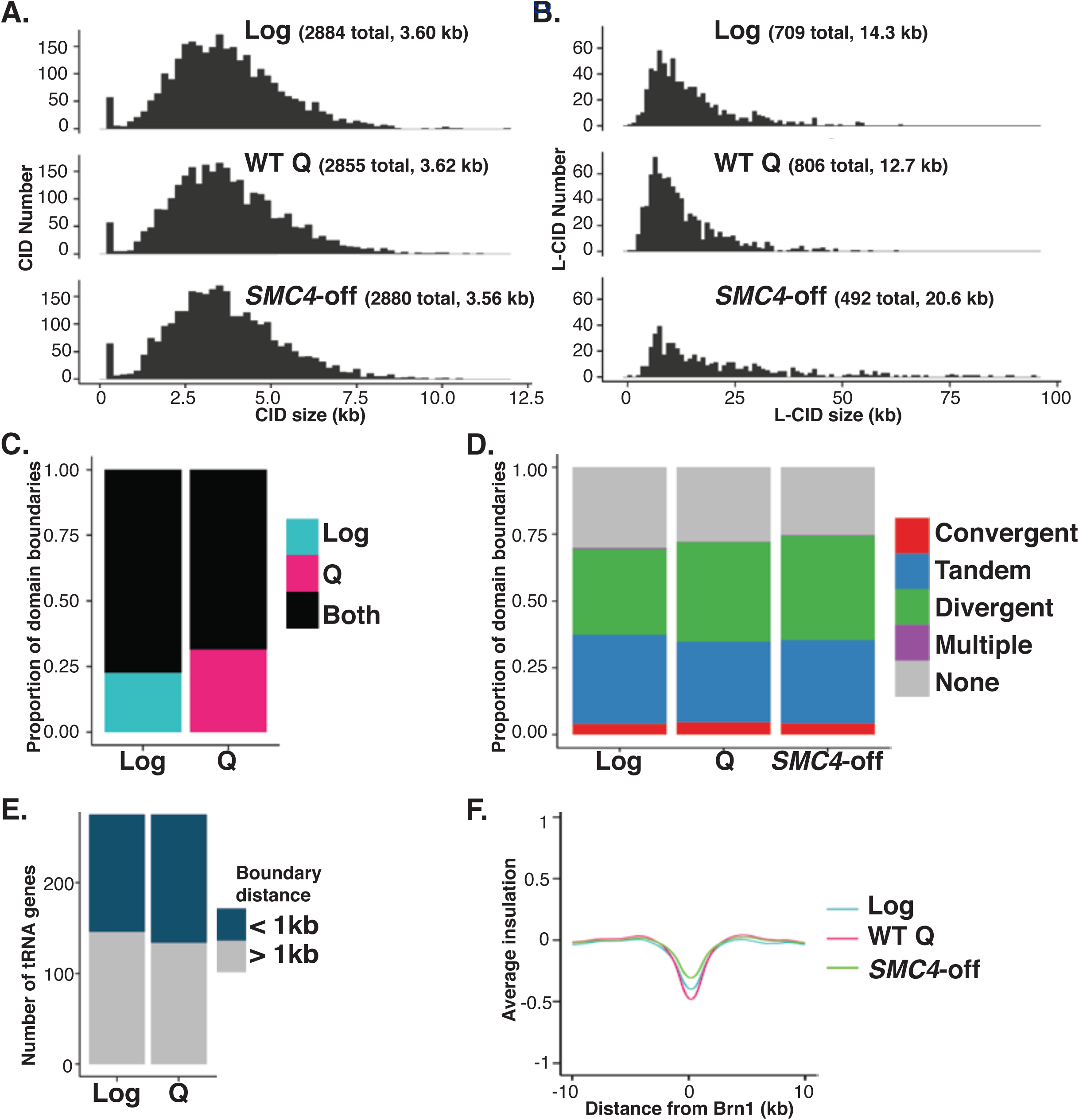
L-CID boundaries are similar in log and quiescence, but L-CIDs are condensin-dependent in quiescence. (**A**) Histogram of the number and size of CIDs in log, WT Q, and *SMC4*-off. Numbers shown are the average number and length of CIDs in each condition. (**B**) Histogram of the number and size of L-CIDs in log, WT Q, and *SMC4*-off. Numbers shown are the average number and length of large domains in each condition. (**C**) Proportion of domain boundaries in log and WT Q that are shared within 1 kb (black) versus unique to log (blue) or Q (red). (**D**) Proportion of L-CID boundaries that overlap with the given class of intergenic region in log, WT Q, and *SMC4*-off. (**E**) Number of tRNA genes within 1 kb of large domain boundaries in log and Q. (**F**) Metaplot of average contact insulation score centered at Brn1 ChIP-seq peaks in Q. Lower scores indicate higher insulation.

**Figure S4.**
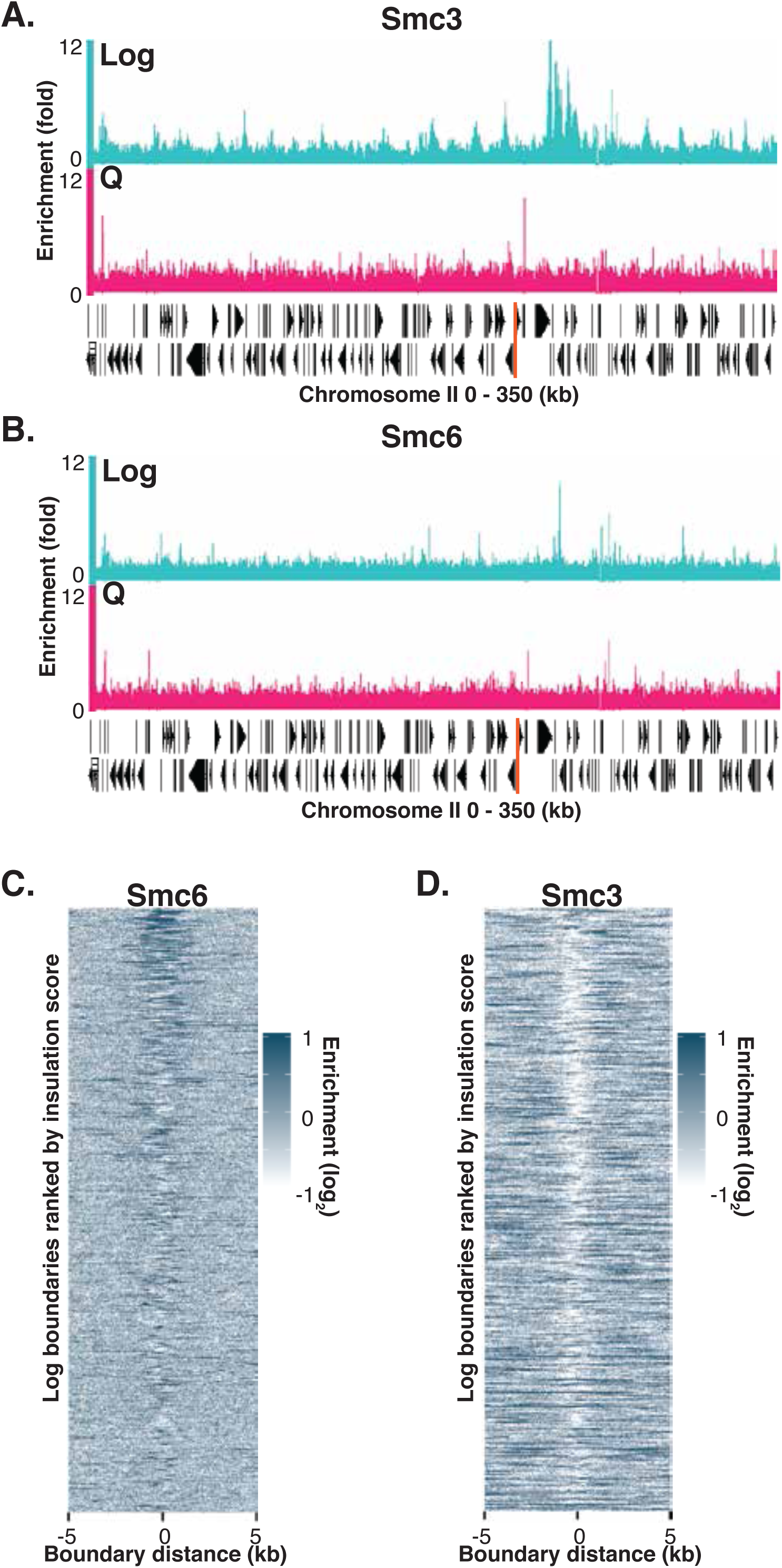
Cohesin and Smc5/6 enrichment is strongly reduced in quiescence. (**A-B**) Example ChIP-seq data of the Smc3 subunit of cohesin (A) and the Smc6 subunit of Smc5/6 (B) in log (blue) and Q (red) across a region of Chromosome II. (**C-D**) Smc6 (C) and Smc3 (D) binding to L-CID boundaries in log.

**Figure S5.**
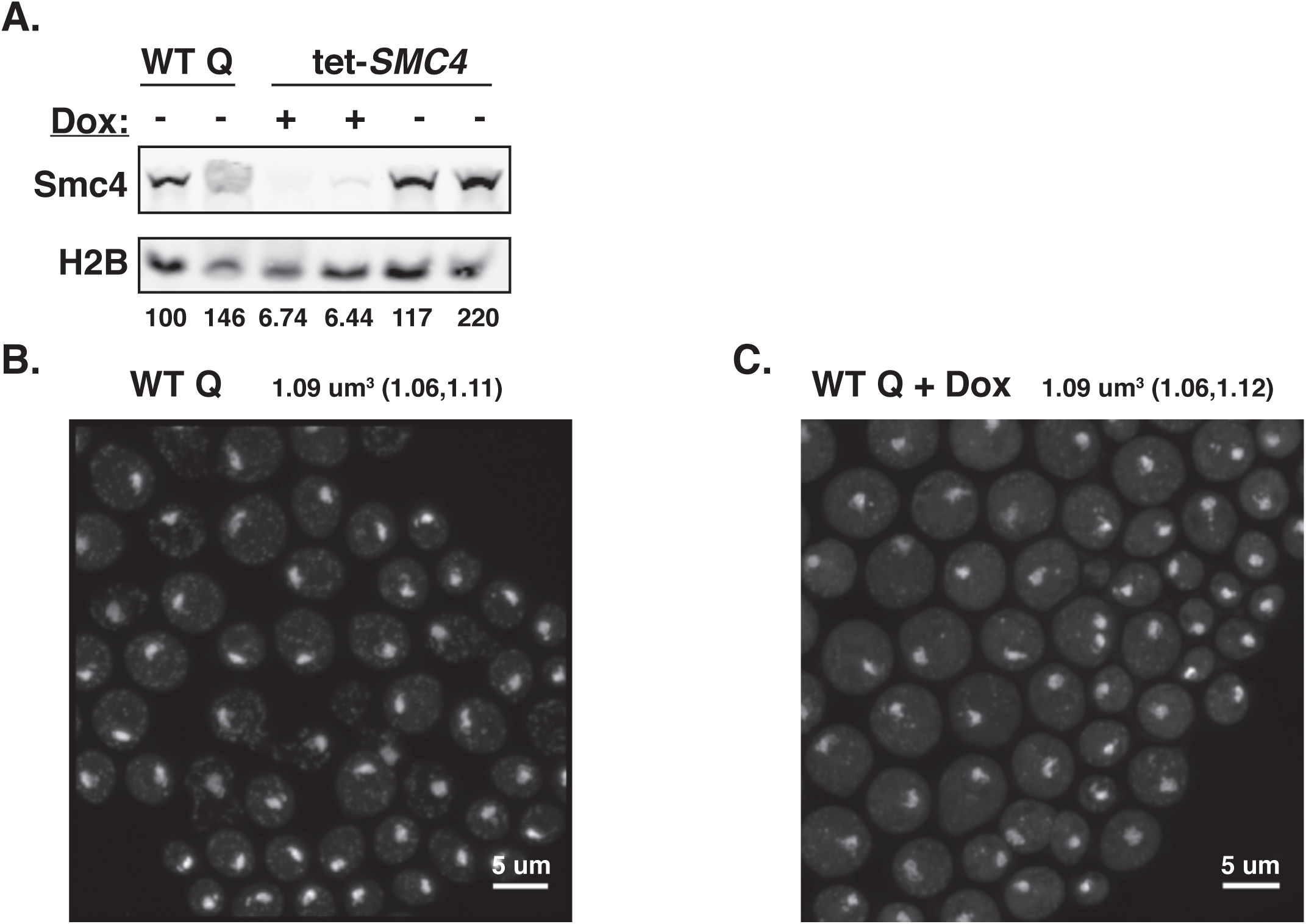
Condensin is depleted in *SMC4*-off cells. (**A**) Western blot of FLAG-tagged Smc4 in WT Q and tet-*SMC4* cells with or without doxycycline addition. Loading control is H2B probed with a polyclonal antibody. Numbers shown are the percentage density of Smc4 bands normalized to H2B versus the normalized WT Q Smc4 band. Band density was quantified using ImageJ software. (**B-C**) DAPI staining and microscopy of purified WT Q cells grown in the presence and absence of doxycycline. Numbers shown are the mean and lower and upper bounds of the 95% confidence interval for the volume of DAPI-stained chromatin.

**Figure S6.**
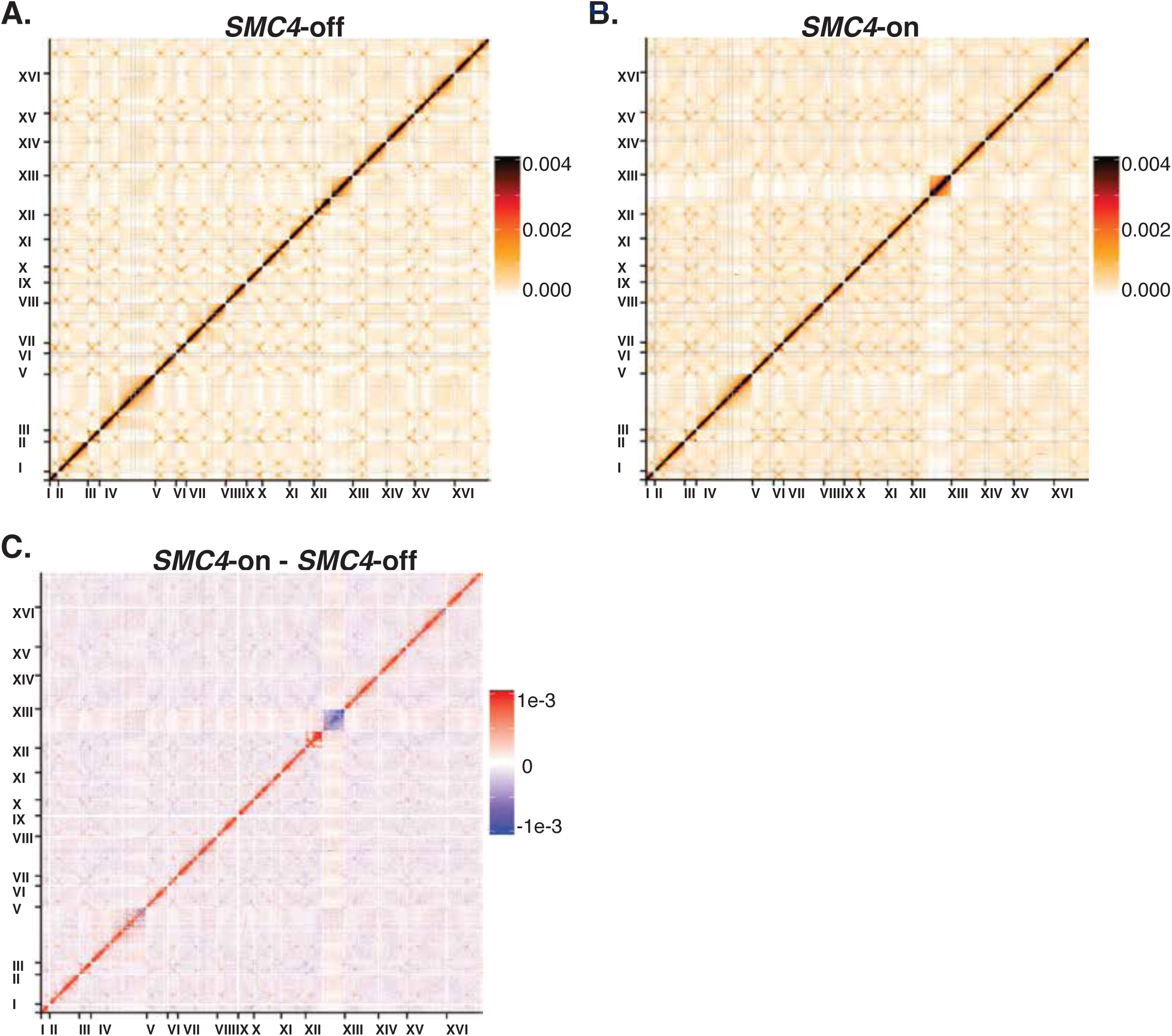
Chromatin conformation changes between *SMC4*-on and *SMC4*-off. (**A**) Genome-wide Micro-C XL data in purified *SMC4*-off Q cells. Each pixel represents normalized contacts within a 5 kb bin. (**B**) Genome-wide Micro-C XL data in purified *SMC4*-on Q cells. (**C**) Difference plot of *SMC4*-off Micro-C data subtracted from *SMC4*-on. Red indicates contacts are higher in *SMC4*-on, blue indicates a greater number of contacts in *SMC4*-off.

**Figure S7.**
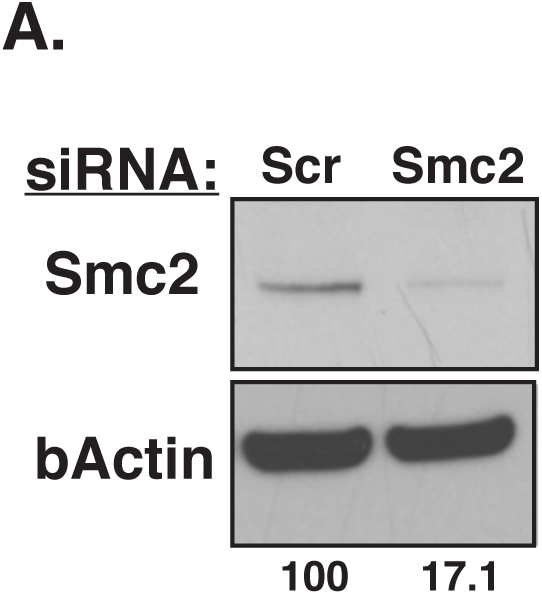
Smc2 knockdown is efficient in quiescent HFFs. (**A**) Western blot of Smc2 and bActin (as a loading control) for cells transfected with siRNA. Numbers shown are the percentage density of the bActin-normalized Smc2 band with siRNA targeting Smc2 (siSmc2) versus the normalized Smc2 band from cells transfected with scrambled control siRNA (Scr). Band density was quantified using ImageJ software.

## Methods

### Yeast Strains, Growth, and Quiescent Cell Purification

All experiments were performed in prototrophic W303 S. *cerevisiae* strains. For ChIP experiments, Brn1, Smc3, and Smc6 were tagged with a C-terminal Myc tag. For the generation of *SMC4-tet* strains, MATa cells harboring a tet-*SMC4*-2L-3FLAG allele and MATa cells harboring a Ssn6-tTA-TetR allele were crossed and dissected. We found that Smc4 depletion during quiescence entry varies among isolates for unknown reasons, so we screened for strains with good Smc4 depletion during quiescence entry by Western blotting. The degree of Smc4 depletion and chromatin decondensation, as observed by DAPI staining, correlated well. For cells arrested in G1, cultures were incubated with alpha-factor at final concentration of 5 ug/mL for 85 minutes at 30°C. For quiescent cell purification (Spain et al., 2018), yeast were grown in rich media for seven days, pelleted, and resuspended in 1 mL of water. To prepare density gradients, 12.5 mL 90% Percoll/150 mM NacL was added to 30 mL round-bottom centrifuge tubes and pre-spun at 10,000 g for 15 min. Yeast were added to the tops of gradients, and gradients were spun at 296 g for 1 hour. Quiescent cells were removed from the bottoms of gradients and quantified via spectrophotometry. For condensin depletion experiments, a stock solution of 20 mg/mL doxycycline resuspended in water was added to cultures to 20 μg/mL approximately 10 hours prior to the diauxic shift, then replenished on days 4 and 7 of culturing prior to quiescent cell purification on day 8. Smc4 depletion was checked for each experiment by Western blot using M2 anti-FLAG antibody (Sigma, catalog #F1804). For thiolutin addition, thiolutin (Tocris, catalog #87-11-6) was resuspended in water to 1 mg/mL, then added to cultures to 3 μg/mL at the diauxic shift and replenished every 12 hours until cell harvesting.

### Yeast Staining and Microscopy

For DAPI staining, 4 optical density units of cells were fixed in 1 ml of 3.7% formaldehyde in 0.1 M KPO_4_ pH 6.4 at 4 C for 20 minutes with rotation. Cells were washed once in 0.1 M KPO_4_ pH 6.4, re-suspended in 1 mL of sorbitol/citrate (1.2 M sorbitol, 100 mM K_2_HPO_4_, 36.4 mM citric acid) and frozen at 20°C. Cells were then digested at 30°C in 200 μl of sorbitol/citrate containing 0.2 μg (for log) or 2.5 μg (for quiescent) of 100T zymolyase for 5 minutes (log) or 30 minutes (quiescent). Cells were washed once and resuspended in sorbitol/citrate, then loaded onto PTFE printed slides from Electron Microscopy Sciences (catalog #63430-04) coated with 0.1% polylysine. After washing twice in sorbitol/citrate, slides were incubated in ice cold methanol for 3 minutes and ice cold acetone for 10 seconds, dried, then washed twice with 10 mg/mL BSA dissolved in PBS. Then 10 μL of DAPI-mount (0.1 μg/mL DAPI, 9.25 mM p-phenylenediamine, dissolved in PBS and 90% glycerol) was added to each slide well. Z-stack images were obtained using a Zeiss 780 NLO confocal microscope with Airyscan at 630X and optimized resolution and nuclear volumes were measured using ImageJ software.

### Chromatin Immunopreciptation

ChIP-seq was performed as previously described (Rodriguez et al., 2014), except using approximately 200 optical density units of log cells and at least 300 optical density units of quiescent cells to prepare 1 mL of cell lysate. Quiescent cells were bead beat for two rounds of 5 minutes each and all cells were sonicated for two rounds of 15 minutes and one round of 10 minutes. The DNA content of cell lysates was measured using the Qubit fluorescence spectrophotometry system, and 1 μg of DNA was used for each ChIP. For H3 ChIP, 1 μl of anti-H3 antibody (Abcam, catalog #1791) was conjugated to 20 μl Dynabeads Protein G beads (Invitrogen, catalog #10004D) per reaction. For Myc ChIP, 5 μl of 9E10 purified anti-Myc (Covance, catalog #139049002) was conjugated to 20 μl Protein G beads. For Pol II ChIP, 1 μl anti-Rpb3 antibody (Biolegend, catalog #665003) was conjugated to 20 μl Dynabeads M-280 sheep anti-mouse IgG beads (Invitrogen, catalog #11201D). Libraries were prepared using the Ovation Ultralow v2 kit (NuGEN, catalog #0344). Single-end sequencing was completed on an Illumina HiSeq 2500 in rapid run mode by the Fred Hutchinson Cancer Research Center genomics core facility.

### ChIP-seq data analysis

For final analysis, fastq files for biological and technical replicates for each sample were merged. Reads were aligned to the sacCer3 reference genome (release R64-2-1) using bowtie2 version 2.2.5 in --very-sensitive mode (Langmead and Salzberg, 2012). Aligned reads were filtered and indexed using SAMtools (Li et al., 2009). Reads were adjusted so that the genome average was set at 1 fold enrichment, then data from immunoprecipitated samples were divided by data from input samples. Peak calling was completed using “callpeak” commands in MACS v2 software (Zhang et al., 2008), and shared peaks between samples were determined using MACS “bdgdiff” commands. Heatmaps were generated using deepTools v2.0 (Ramirez et al., 2016). The most and least activated genes were calculated using a ratio of the average Pol II enrichment over the length of each open reading frame in WT versus *SMC4*-off.

### Micro-C XL

Micro-C XL was performed as described using pelleted chromatin except with the following modifications (Hsieh et al., 2016). For log, two cultures of 55 optical density cells each were used so that the entire Micro-C protocol was performed in two separate reactions through the end repair and ligation steps, then DNA from extracted bands during the gel purification step was combined into a single reaction for the duration of the protocol. The cell walls of log cells were permeabilized using 250 μl of 10 mg/mL 20T zymolyase, shaking for 30 minutes at 30°C. For quiescent cells, eight reactions of 200 optical density units of purified cells each were used so that the entire Micro-C protocol was performed in eight separate reactions through the end repair and ligation steps. DNA was combined into two reactions during the phenol-chloroform and ethanol precipitation steps and loaded onto the purification gel in two lanes. DNA from extracted bands was combined into a single reaction for the duration of the protocol. The cell walls of quiescent cells were permeabilized using 1 mL of 10 mg/mL 100T zymolyase with shaking at 30°C for approximately 2 hours, or until over 50 percent of cells appeared lysed under a microscope. For each preparation of log and quiescent cells, an additional two reactions’ worth of cells (2 x 55 optical densities for log and 2 x 200 optical densities for quiescence) were split into four reactions and titrated with MNase to determine the appropriate concentration of MNase for the experiment. The concentration of MNase used was determined to be the concentration that gave approximately 90 percent mononucleosome-sized fragments in the pelleted chromatin sample. Paired-end sequencing was completed on an Illumina HiSeq 2500 in rapid run mode by the Fred Hutchinson Cancer Research Center genomics core facility.

### Micro-C XL data analysis

Reads were mapped independently to the sacCer3 reference genome (release R64-2-1) using bowtie2 version 2.2.5 with the --very-sensitive parameter set (Langmead and Salzberg, 2012). Except for rDNA interaction analyses, all read pairs where both reads mapped with MAPQ >= 6 were deduplicated, and filtered to exclude in-facing read pairs with ends < 400 bp apart, which likely represent unligated fragments. The resulting read pairs were ICE normalized (Imakaev et al.) using the cooler suite (https://github.com/mirnylab/cooler) balance function, with settings --ignore-diags 1 --mad-max 9. For rDNA interaction analyses, all read pairs where one of the two reads mapped to the region chrXII:451575-468900 were mapped to genomic bins, normalized by the ICE normalization weights for the full genomic matrix, and then divided by the average across all bins. *P(s)* genomic distance decay curves (Naumova et al.) were calculated by dividing the number of interactions in 1.03-fold genomic distance bins by the total number of possible interactions at each distance, and then normalizing so that the sum across all bins equals 1. For reproducibility analyses, the hicrep R package (Yang et al., 2017) was run on all pairs of experiments individually subsampled to 6,000,000 unique contacts (read pairs excluding unligated fragments), at 5 kb resolution for up to 100 kb interactions. The median stratum-adjusted correlation coefficient across the 16 chromosomes was calculated. Based on the similarity of replicate experiments, all replicates were merged for remaining analyses. CID boundaries were called using the cworld-dekker package (https://github.com/dekkerlab/cworld-dekker) (Giorgetti et al., 2016) matrix2insulation.pl script with settings --is2400 --nt0 --ids1600 --ss400 --im mean. L-CID boundaries were called using matrix2insulation.pl with settings --is4800 -- nt0.4 --ids3200 --ss800 --im mean. CIDs and L-CIDs were called using matrix2tads.pl with the corresponding insulation score tracks using default settings.

### Human Cell Culture Growth and Microscopy

Human foreskin fibroblasts (HFFs) were maintained in DMEM supplemented with 10% FBS (Peak Serum) and 1% penicillin/streptomycin (Life Technologies). For proliferative conditions, cells were plated into an 8-well chamber slide (Fisher) and cultured in growth media (10% FBS) for 48 h before fixation. For quiescent conditions, cells were transfected with 20 nM scrambled (Allstars negative control siRNA, Qiagen) or human SMC2-specific siRNAs (Qiagen) using Lipofectamine RNAiMAX (Life Technologies), according to the manufacturer’s instructions. 24 h post-transfection, cells were plated into 8-well chamber slides or 35 mm tissue culture dishes (USA Scientific) in growth media. 48 h later, cells were switched to DMEM with 0.1% FBS for 4 days to induce quiescence.

For immunofluorescence, cells were fixed with 4% formaldehyde for 15 min, washed with 1xPBS, and permeabilized with 0.1% Triton X-100 for 10 min at room temperature. Cells were then blocked with DAKO protein block (Agilent) for 1 h, followed by overnight incubation with anti-Ki-67 antibody (1:250, ab16667, Abcam) or anti-Cyclin D1 antibody (1:100, #2978, Cell Signaling) in DAKO antibody diluent (Agilent) at 4°C. The next day, cells were washed with 1 x PBS and incubated with secondary antibody conjugated with AlexaFluor 568 (1:400, Life Technologies) in DAKO antibody diluent for 1 h at room temperature. Slides were mounted with Prolong Gold antifade reagent with DAPI (Life Technologies).

Images were acquired using a DeltaVision Elite imaging system (GE) with an Olympus IX71 inverted microscope and a 100x/1.4 UPlanSApo objective. Images were deconvolved using Softworx 3.4.3, and Z projections were generated using five consecutive images taken at an interval of 0.2 μm. To analyze DAPI intensity, a threshold value was set, and the fraction of area in each nucleus with intensity above the threshold was measured using ImageJ. Only Cyclin D1-positive (indicating G1) cells and Ki-67-negative (indicating G0) cells were analyzed. Significant differences in area fractions were calculated using t-test.

For immunoblot, cells were lysed in RIPA buffer, and the protein concentrations were determined by the BCA assay (Pierce). 15 μg of each lysate was loaded onto a NuPAGE Bis-Tris Protein Gel (Life Technologies) and transferred to nitrocellulose for blocking and incubation with primary antibody and secondary antibody conjugated with HRP for detection by ECL. Antibodies were obtained from the following sources: anti-SMC2 (A300-058A) from Bethyl, anti-p21 (2947), anti-p27 (3686), anti-rabbit IgG (7074) and anti-mouse IgG (7076) linked with HRP from Cell Signaling, and anti-bActin (A5441) from Sigma.

